# Integrated Analysis of Single-Cell and Bulk RNA-Seq Data reveals that Ferroptosis-Related Genes Mediated the Tumor Microenvironment predicts Prognosis, and guides Drug Selection in Triple-Negative Breast Cancer

**DOI:** 10.1101/2024.07.04.602021

**Authors:** Xuantong Gong, Lishuang Gu, Di Yang, Yu He, Qian Li, Hao Qin, Yong Wang

## Abstract

**Background:** Triple-negative breast cancer (TNBC) is aggressive, lacking methods to predict recurrence and drug sensitivity. Ferroptotic heterogeneity varies in TNBC subtypes. However, the tumor microenvironment (TME) mediated by ferroptosis genes is unclear. Our study aims to integrate single-cell and bulk RNA sequencing (RNA-seq) data to reveal the ferroptosis-mediated TME in TNBC, predicting prognosis and guiding treatment.

**Methods:** The single-cell and bulk RNA-seq data of TNBC were sourced from the Gene Expression Omnibus (GEO) database. Using these data, a machine learning algorithm was employed to integrate and analyze the characteristics of the TME mediated by ferroptosis-related genes in TNBC. Prediction models for TNBC survival prognosis and drug treatment response were established and then validated in an independent set.

**Results:** At the individual cell level, T cells were categorized into three distinct subpopulations, and local macrophages into two subpopulations. The infiltration degree of these different cell subpopulations was closely associated with prognosis and treatment outcomes. Based on this, the risk score model we developed effectively predicted recurrence-free survival in TNBC patients, with independently validated pooled predicted 3-, 4-, and 5-year Area Under the Curves (AUCs) of 0.65, 0.67, and 0.71, respectively. Additionally, we found that patients in the high-risk group may be more responsive to 27 drugs.

**Conclusions:** We have uncovered the tumor immune cell clusters in TNBC mediated by ferroptosis. A risk score model was constructed to identify high-risk TNBC patients, which can assist physicians in disease monitoring and precision therapy. The genes identified hold significant potential as therapeutic targets for TNBC patients.

**Funding:** This project is funded by the National Natural Science Foundation of China (81974268, 82472000, 82304151), the Talent Incentive Program of Cancer Hospital Chinese, Academy of Medical Sciences (801032247), the Cooperation Fund of CHCAMS (CFA202202023), and the open project of Beijing Key Laboratory of Tumor Invasion and Metastasis Mechanism, Capital Medical University(2023ZLKF03).

**Impact Statement:** Integrating single-cell and bulk RNA-seq data elucidates the role of ferroptosis in TNBC, offering a prognostic model and personalized therapeutic insights.

## 1. Background

Global cancer statistics show that breast cancer is the most common cancer in women and the leading cause of cancer deaths ^[1]^. Triple-negative breast cancer (TNBC) is a subtype of breast cancer that lacks expression of estrogen receptor, progesterone receptor and human epidermal growth factor receptor-2. Compared with other subtypes, TNBC is highly aggressive, with a poor overall prognosis for patients and a median survival of less than 1 year after recurrent metastasis ^[2–5]^. Due to the lack of targets for endocrine therapy and targeted therapeutic agents, chemotherapy and emerging immunotherapy are the main therapies. TNBC is highly heterogeneous and only some patients benefit from treatment ^[6–8]^. Unfortunately, there is no effective method that can effectively predict the prognostic risk and drug sensitivity of TNBC, which is an urgent problem in the clinic.

The tumor microenvironment (TME) is composed of various cell types such as cancer-associated fibroblasts, tumor-associated macrophages, T cells, NK cells, B cells, endothelial cells, etc., which interact with cancer cells and influence various aspects such as tumor progression, metastasis, and response to therapy ^[9–11]^. The composition and functional status of TME varies greatly among different patients with breast cancer, and revealing the TME of each patient is crucial for selecting a reasonable treatment and controlling tumor progression in the long term ^[12–13]^.

Ferroptosis is an iron-dependent form of non-apoptotic, oxidative form of regulated cell death involving lipid hydroperoxides ^[14–15]^. Studies have shown that drug-resistant cancer cells are more sensitive to ferroptosis ^[16–18]^. Therefore, ferroptosis is more recognized as a potential target for cancer therapy. Ferroptotic heterogeneity exists in different subtypes of TNBC ^[19]^. Since ferroptosis is regulated by multiple metabolic pathways, the ferroptosis landscape of TNBC remains unexplored and its relationship with patient prognosis and treatment response is uncertain ^[20]^. Therefore, exploring ferroptosis-related genes in TNBC and their correlation with the immune microenvironment may provide a new treatment trend for TNBC.

RNA sequencing (RNA-seq) technology is a gene expression analysis method that can qualitatively and quantitatively explore the transcriptome characteristics of biological samples at the tissue and cellular levels ^[21–22]^. Single-cell RNA-seq (scRNA-seq) has greatly enhanced our understanding of transcriptional heterogeneity across cell types and states, and can be used to explore the TME ^[23–24]^. Bulk RNA-seq is used to analyze the average expression levels of RNA in tissues or cell populations, and can be utilized to explore differences among individuals ^[25–26]^. Machine learning is a branch of artificial intelligence that enables computer systems to learn from training data and gain experience, and it is increasingly being used for the diagnosis, treatment, and prognostic evaluation of cancer ^[27–28]^. Literature reports that several models have been constructed based on RNA-seq data for the prognostic evaluation of breast cancer patients, but the reproducibility and interpretability of the results still deserve further exploration ^[29–32]^. This study intends to further explore the relationship among ferroptosis genes, prognosis, and drug sensitivity from the perspective of ferroptotic heterogeneity in TNBC. In this work, we collected and integrated data from single-cell and bulk RNA-seq to characterize the ferroptosis-related gene mediated TME landscape of TNBC on multiple scales. furthermore, based on the findings in TME, predictive models of patient prognosis and treatment response were constructed using machine learning algorithms. We hope to provide a new way of individualized risk management for triple-negative breast cancer patients and assist their precision treatment.

## 2. Methods

| Key Resources Table |  |  |  |  |
| --- | --- | --- | --- | --- |
| Reagent type (species) or resource | Designation | Source or reference | Identifiers | Additional information |
| software,<br>algorithm | R package<br>Seurat | Microsoft | N/A | Version 4.1.1 |

### 2.1 Patient data collection and processing

The data used in this study were collected from public datasets. Nine single-cell RNA-seq TNBC samples were obtained from the Gene Expression Omnibus (GEO) database (https://www.ncbi.nlm.nih.gov/geo/query/acc.cgi) under the accession number GSE176078 ^[33]^ (Table1).

**Table 1.**
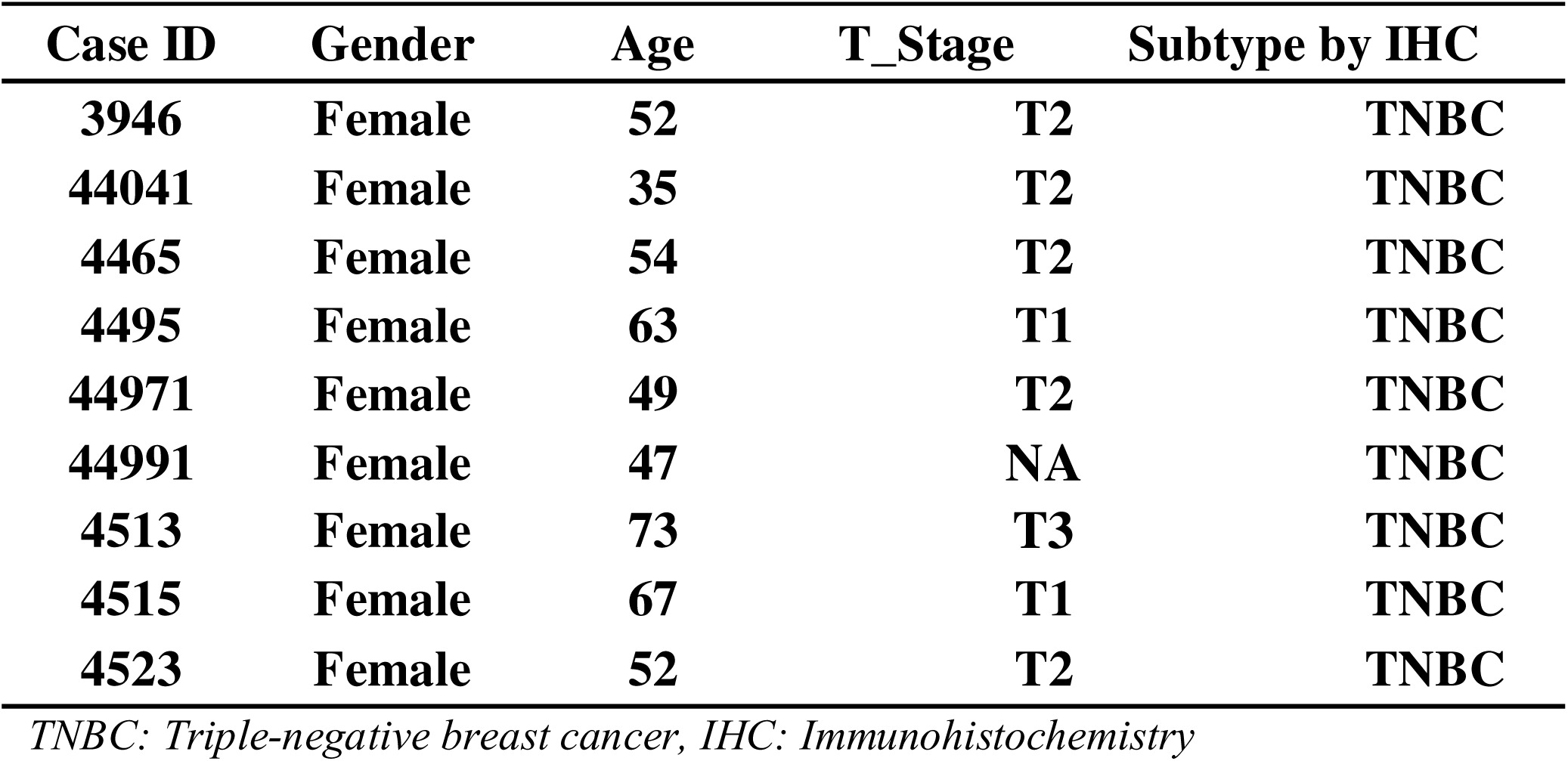
Sample information for triple-negative breast cancer in GSE176078.

To construct the model to predict prognosis and immunotherapy efficacy, bulk RNA-seq datasets and clinicopathological information of TNBC were obtained from the GEO database under the accession number GSE25066 ^[34]^. The data for external validation were obtained from the GEO database under the accession number GSE86166 ^[35]^.

### 2.2 Single-cell RNA-seq data processing

Data were processed using the R package Seurat (version 4.1.1) for quality control (QC). Based on QC metrics suggested in Scanpy tutorial, outlier cells were removed based on relevant feature data (nFeature_RNA, nCount_RNA, percent.mt), i.e., cells with less than 200 genes expressed or more than 20% mitochondrial genes counts were filtered out. After quality control, the expression data of each cell was normalized separately using the NormalizeData function and the top 2000 highly variable genes (HVGs) of each cell were identified using the FindVariableFeatures function. To remove batch effects among different samples, we utilized the FindIntegrationAnchors function to find anchor genes and then the reclassified datasets were integrated utilized the IntegrateData function. Data normalization was performed using the ScaleData function in Seurat. The RunPCA function was utilized to reduce the dimension of principal component analysis (PCA) for the first 2000 HVGs screened above.

### 2.3 Genes associated with ferroptosis acquiring

The data of 471 genes (Supplementary Table 1) associated with ferroptosis were collected from the FerrDb website (http://www.zhounan.org/ferrdb/current/). Then, from the above genes, screened for genes with expression information in the single-cell matrix of TNBC patients. Firstly, based on the clinical data of the samples, the expression values of ferroptosis-related genes in the same type of samples were averaged and presented in a heatmap. Afterwards, the expression values of ferroptosis-related genes were averaged across different cell types and also presented in a heatmap.

### 2.4 Cell subtype identification associated with ferroptosis

Because information of cell type annotation was listed in the original literature of the GSE176078 dataset, the metadata of the original literature was used directly for the next analyses. The R package Seurat (version 4.1.1) was used to identify cell types and presented results via t-SNE. Non-negative matrix decomposition (NMF) is an algorithm based on high-throughput data to identify and cluster out different molecular functional patterns ^[36]^. In this work, subtypes of macrophages and T cells based on genes associated with ferroptosis using the “NMF” R package (v 0.26).

### 2.5 Cell-cell communication analysis

Cell-cell communication mediated by ligand-receptor complexes plays an important role in multiple biological processes ^[37]^. In this study, the “iTALK” R package was utilized to construct the cellular communication network. First, use the rawParse function to identify the top 50% highly expressed genes for each cell type based on their expression means. Then, utilize the FindLR function to identify ligands and receptors among the highly expressed genes. Finally, construct an interaction network based on all the interaction relationships.

### 2.6 Pseudotime analysis

Pseudotime analysis of scRNA-seq snapshot data helps to provide an approximate landscape of gene expression dynamics ^[38]^. The “Monocle” R package (v 2.28.0) was applied for pseudotime analysis to conduct cellular trajectory. The reduceDimensio function based on the DDRTree algorithm is used to reduce the dimensions of the data. Based on the Progenitor Cell Biology Consortium (PCBC) database (https://www.synapse.org), the stemness signature was identified via the one-class logistic regression (OCLR) algorithm, then the stemness index of each TNBC cell was calculated by scaling the Spearman correlation coefficients to be between 0 and 1. Eventually, the order Cells function was utilized to sort cells and complete construction of trajectory.

### 2.7 Transcriptional factor analysis

The Single-cell Regulatory Network Inference and Clustering tool (SCENIC) enables simultaneous gene regulatory network reconstruction and cell-state identification from scRNA-seq data ^[39–40]^. We utilized the “SCENIC” R package (v1.3.1) to establish the transcription factor (TF) regulatory network. Specifically, first, coexpression modules are inferred using the “GENIE3/GRNBOOST” function to identify gene set with co-expressed TFs. Next, the indirect targets are pruned from these modules using cis-regulatory motif discovery (cisTarget). Finally, the AUCell algorithm was utilized to evaluate the activity of regulons.

### 2.8 Hallmarks gene set enrichment analysis

Hallmarks gene set enrichment analysis was utilized the “irGSEA” R package (v2.1.5). The ssGSEA algorithm was utilized to conduct differential pathway score between ferroptosis-related immune cell subtypes.

### 2.9 Construction and validation of the prognostic model

First, based on the recurrence-free survival (RFS) data of 178 TNBC patients in the GSE25066 database, this study utilized the “survival” (v3.2-7) and “survminer” (v0.4.8) R package to perform univariate Cox proportional hazards regression analyses on ferroptosis-related genes signature of different immune cell subtypes.

The genes screened for significant association with RFS in the regression analysis (p<0.01) were utilized to construct a risk factor-based model by the Least Absolute Shrinkage and Selection Operator (LASSO) method implemented in the “glmnet” R package (v4.0-2). TNBC patients were divided into low- and high-risk groups according to the median risk score, and Kaplan-Meier survival curves were utilized to compare the RFS rates between the two groups, p value < 0.05 was considered to significance. To validate the performance of the model, receiver operating characteristic (ROC) curves were demonstrated and the AUC values were calculated for evaluating 3-year, 4-year, and 5-year RFS rates of TNBC patients in GSE25066 database. In addition, the GSE86166 as well as the TCGA database were utilized as external validation sets to further verify the robustness of the performance of the constructed model in this study.

### 2.10 Immune cells infiltration and drug sensitivity analysis

Immune cells infiltration is important to the anti-tumor response, and these cells are diverse among patients ^[41–42]^. The “CIBERSORT”, “GSVA” and “TIMER” R package was utilized to determine the distribution of different immune cell types between low- and high-risk groups. Next, the 50% inhibitory concentration (IC50) of 138 chemotherapeutic drugs was calculated for each patient using the “pRRophetic” R package (v 0.5), and drugs significantly associated with the risk score were screened. Finally, we calculated the tumor immune dysfunction and exclusion (TIDE) score (http://tide.dfci.harvard.edu.) for each patient and thus analyzed its difference between low- and high-risk groups.

### 2.11 Statistical analysis

The R software (version 4.1.3) was utilized for all data analysis. Wilcoxon-rank sum test was utilized to analyze associations of continuous variables. Log-rank test was utilized to analyze differences in survival curves between groups. P value less than 0.05 was considered statistically significant.

## 3. Results

### 3.1 The single-cell landscape of TNBC samples

In this study, a total of 9 TNBC single-cell samples were included. After QC, 38985 genes from 29733 cells per sample were finally selected for subsequent analysis (Supplementary Figure 1). Based on original annotations of data, these cells were divided into 9 major cell clusters and 29 minor cell clusters (Figure 1a-c). Major cell clusters include B-cells, Cancer-associated Fibroblasts (CAFs), Cancer Epithelial, Endothelial, Myeloid, Normal Epithelial, Plasmablasts, Perivascular-like Cells (PVL), T-cells. Of these, Myeloid cluster is further divided into Cycling_Myeloid, Dendritic Cells (DCs), Macrophage, and Monocyte clusters. As shown in Figure 1c, the immune microenvironment of TNBC is characterized by a high proportion of T cells and macrophages. On this basis, ligand-receptor interactions between different major cell clusters are frequent (Figure 1d).

**Fig. 1.**
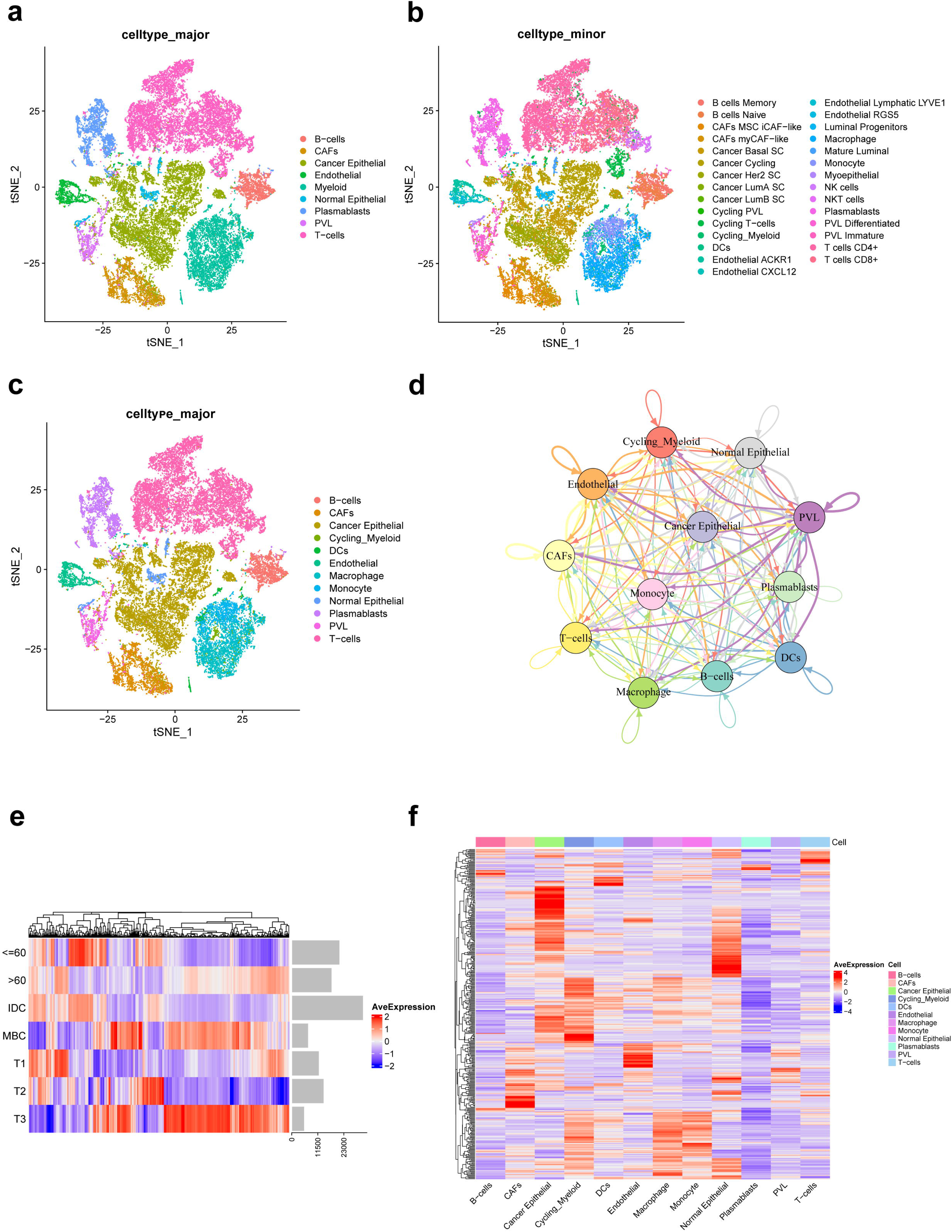
Integration and clustering of scRNA-Seq data from triple-negative breast cancer. (a) t-SNE plot of the 9 major cell clusters. (b) t-SNE plot of the 29 minor cell clusters. (c) t-SNE plot of the 12 major cell clusters. (d) The number of ligand-receptor interactions of different major cell clusters in cell-cell communication network, different colors represent different cell clusters and arrows represent ligand-receptor orientation. (e) Heat map showing the average expression of ferroptosis-related genes in different clinicopathological classifications. (f) Heat map showing the average expression of ferroptosis-related genes in different major cell clusters.

Of the 471 ferroptosis-related genes from the FerrDb website, 391 were present in the single-cell expression matrix of TNBC. We found that there were significant differences in the average expression of these genes in different clinicopathological classifications, as shown in Figure 1e. Moreover, there were significant differences in the expression of ferroptosis-related genes in 12 major cell clusters of TNBC (Figure 1f).

### 3.2 Ferroptosis-related subpopulations of T cells in the TNBC

A total of 11,784 T cells in the scRNA-seq data of this study, including cycling T cells, NK cells, NKT cells, T cells CD4+, and T cells CD8+ identified clusters (Figure 2a). Based on the NMF algorithm (rank=3), each T cell cluster was further categorized into 3 ferroptosis-related subpopulations (T_C1, T_C2, T_C3). Figure 2b shows that differentially expressed top 50 ferroptosis-related genes differed significantly among these 3 subpopulations. Among them, the proportion of NK cells in the T_C2 subpopulation was significantly more than the other two subpopulations (Figure 2c). According to pseudotime trajectories, we found that all three T cell subpopulations were involved at different periods of differentiation (Figure 2d-f). Further, Figure 2g shows the average expression of signature genes associated with 8 functions in 15 subpopulations of T-cells, and we found that gene expression was significantly higher in the Cycling T-cells. Moreover, different T-cell subpopulations play an important role in the anti-tumor process (Figure 2h).

**Fig. 2.**
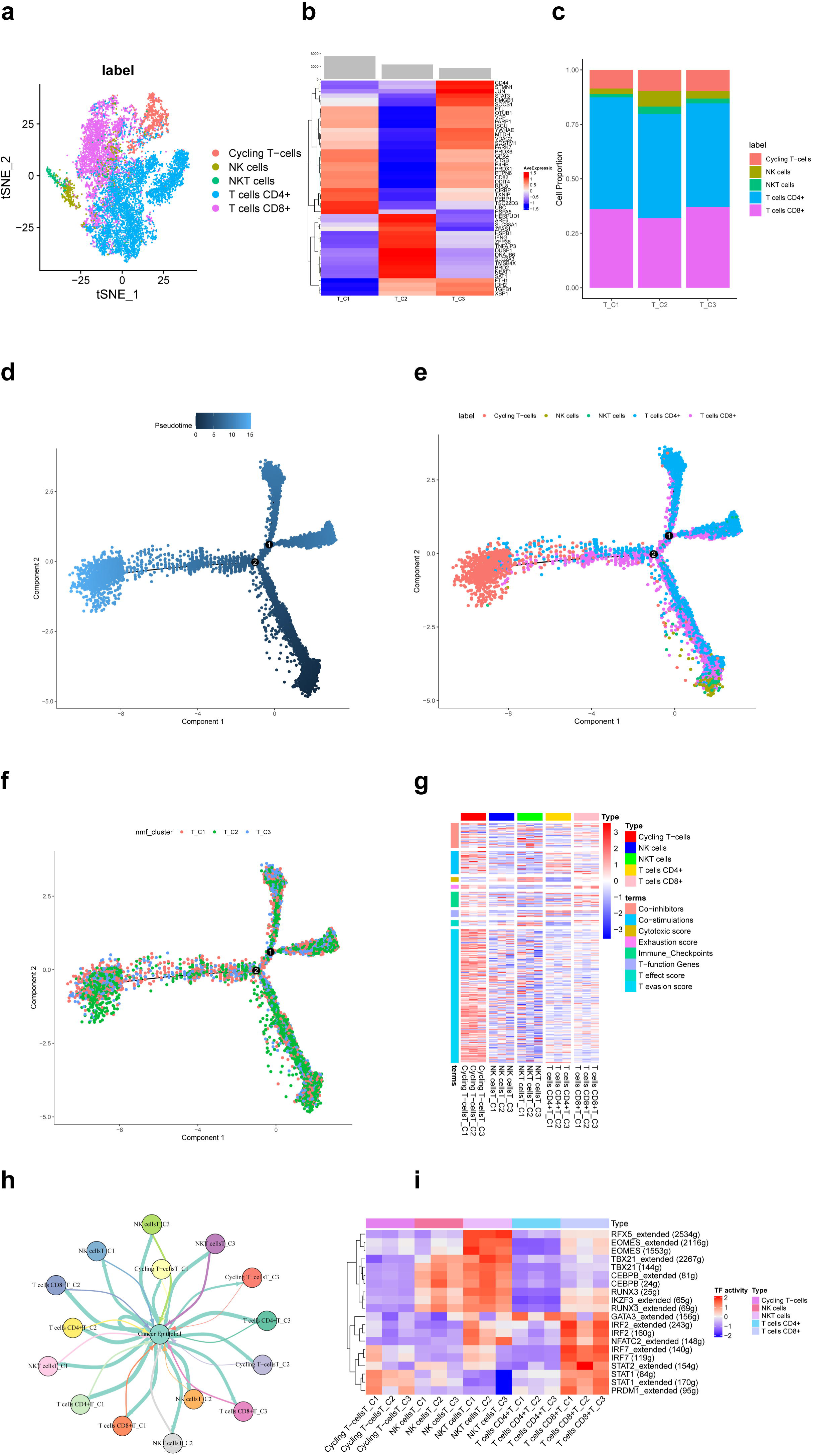
Ferroptosis-related subpopulations of T cells in triple-negative breast cancer. (a) t-SNE plot of the 5 identified clusters of T cells. (b) Heat map showing the expression of ferroptosis-related genes (Top 50) in 3 subpopulations of T cells. (c) Demonstration of the proportion of clusters of T cells in ferroptosis-related subpopulations. (d) Pseudotime trajectories showing the developmental time course of T cells. (e) Pseudotime trajectories of the 5 identified clusters of T cells. (f) Pseudotime trajectories of ferroptosis-related subpopulations of T cells. (g) Heat map showing the average expression of signature genes associated with 8 functions in 15 subpopulations of T-cells. (h) Network diagram of the cell-cell communication between cancer epithelial cells and T cells. (i) Heat map for differential analysis of transcription factors activity.

TFs are key regulators in cellular signal transduction ^[43]^. As a result, RFX5, EOMES, TBX21, CEBPB, RUNX3 and IKZF3 showed higher activities in NKT cells. Besides, IRF, NFATC2, STAT and PRDM1 showed higher activities in CD8^+^ T cells (Figure 2i). The HALLMARK analysis results revealed a prominent enrichment in pathways such as pathways in DNA−Repair and adipogenesis in cycling T cells, whereas in other clusters of T cells, pathways such as epithelial-mesenchymal transition and KRAS signaling had higher enrichment. However, the difference in enrichment scores between ferroptosis-related subpopulations of T cells was not significant.

### 3.3 Ferroptosis-related subpopulations of macrophages in the TNBC

A total of 3671 macrophages in the scRNA-seq data of this study. Based on the NMF algorithm (rank=4), 2 ferroptosis-related subpopulations of macrophages (M_C1, M_C2) were finally obtained after clustering cell clusters with similar markers, the t-SNE plot was shown in Figure 3a. The pseudotime trajectories showed that M_C2 cells belong to the early stage of macrophages and then gradually develop into M_C1 cells (Figure 3b-c). Figure 3d shows that differentially expressed top 50 ferroptosis-related genes differed significantly among subpopulations.

**Fig. 3.**
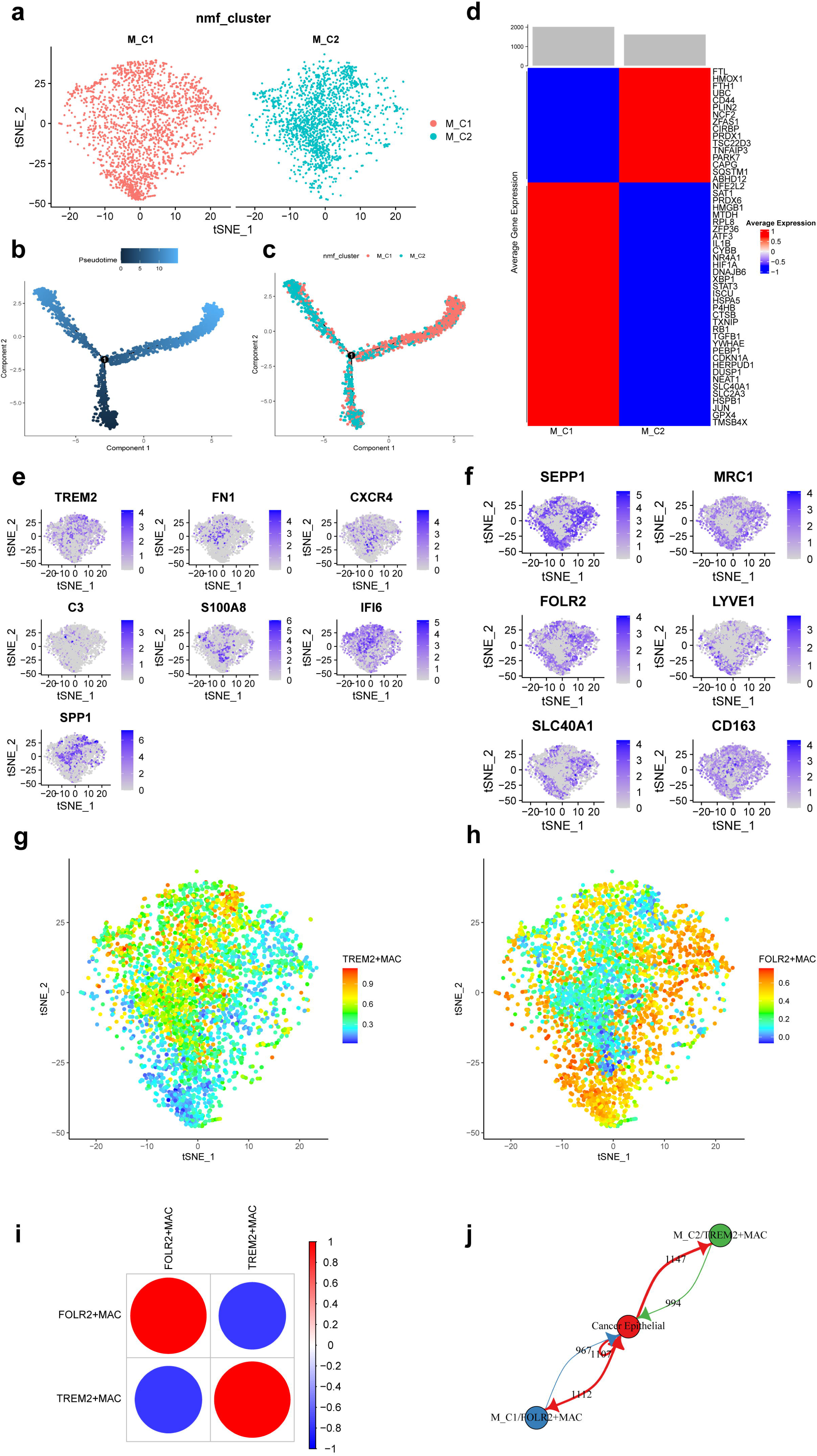
Ferroptosis-related subpopulations of macrophages in triple-negative breast cancer. (a) t-SNE plot of 2 ferroptosis-related subpopulations of macrophages. (b) Pseudotime trajectories showing the developmental time course of macrophages. (c) Pseudotime trajectories of 2 ferroptosis-related subpopulations of macrophages. (d) Heat map showing the average expression of ferroptosis-related genes in 2 subpopulations of macrophages. (e) t-SNE plot showing the expression patterns of marker genes in M_C2. (f) t-SNE plot showing the expression patterns of marker genes in M_C1. (g) t-SNE plot showing the enrichment score of the TREM2+MAC in each cell. (h) t-SNE plot showing the enrichment score of the FOLR2+MAC in each cell. (i) Bubble plot showing the correlation of enrichment score between TREM2+MAC and FOLR2+MAC. (j) Network diagram of the cell-cell communication between cancer epithelial cells and ferroptosis-related subpopulations of macrophages.

Further, we found that FOLR2, SEPP1, MRC1, LYVE1, SLC40A1, and CD163 were highly expressed in M_C1 cells and named M_C1 as FOLR2+MAC. whereas, TREM2, FN1, CXCR4, C3, S100A8, IFI6, and SPP1 were highly expressed in M_C2 and named M_C2 as TREM2+MAC. Figure 3e-f shows the marker genes in each subpopulation of macrophages. Enrichment scores of the 2 subpopulations in each cell are shown by t-SNE plots (Figure 3g-h). Moreover, the enrichment score of TREM2+MAC and FOLR2+MAC were significantly negatively correlated (r=-0.779) (Figure 3i). Communication between these two subpopulations and cancer epithelial cells is vigorous (Figure 3j). In addition, detection of TFs activity in each macrophage cell revealed that most of TFs had higher activity in FOLR2+ MAC cells (Supplementary Figure 2).

### 3.4 Construction and validation of predictive survival models based on ferroptosis-related genes

Based on the RFS data of TNBC in the GSE25066 database, univariate Cox regression analysis was performed on 8371 specific markers of ferroptosis-related subpopulations, and 41 genes significantly associated with RFS were screened (p < 0.01) (Supplementary Table 2, Supplementary Figure 3). These genes were further analyzed by Least Absolute Shrinkage and Selection Operator (LASSO) regression, and 23 genes were finally selected for the construction of the risk model. We calculate the risk score using the following formula: risk score = TMEM160 * (−0.4689) + EWSR1 * (−0.4237) + BCAT2 * (−0.2346) + PNKP * (−0.1592) + MLEC * (−0.1567) + SAP30BP * (−0.1469) + TBL1XR1 * (−0.0894) + STAG1 * (−0.0695) + NR4A1 * (−0.0630) + TNFRSF9 + (−0.0503) + PSD3 * (−0.0153) + BAD * (−0.0094) + MIIP * 0.0104 + HIST3H2A * 0.0294 + CDC25B * 0.0528 + TCEB1 * 0.0670 + HMGCS1 * 0.0988 + SPC25 * 0.1317 + TKT * 0.2234 + PTTG1 * 0.2960 + ADA * 0.3097 + AK1 * 0.4003 + AIMP2 * 0.4081 (Figure 4a-c). All patients were categorized into low- and high-risk groups according to the median value of the risk score (Figure 4d-f). Survival curves showed that patients in the high-risk group had a shorter RFS compared to patients in the low-risk group (p<0.05, Figure 4g). In addition, the risk score had an excellent predictive effect in predicting RFS in TNBC patients, its AUCs for 3-, 4-, and 5 years RFS were 0.87, 0.88, and 0.88 (Figure 4h).

**Fig. 4.**
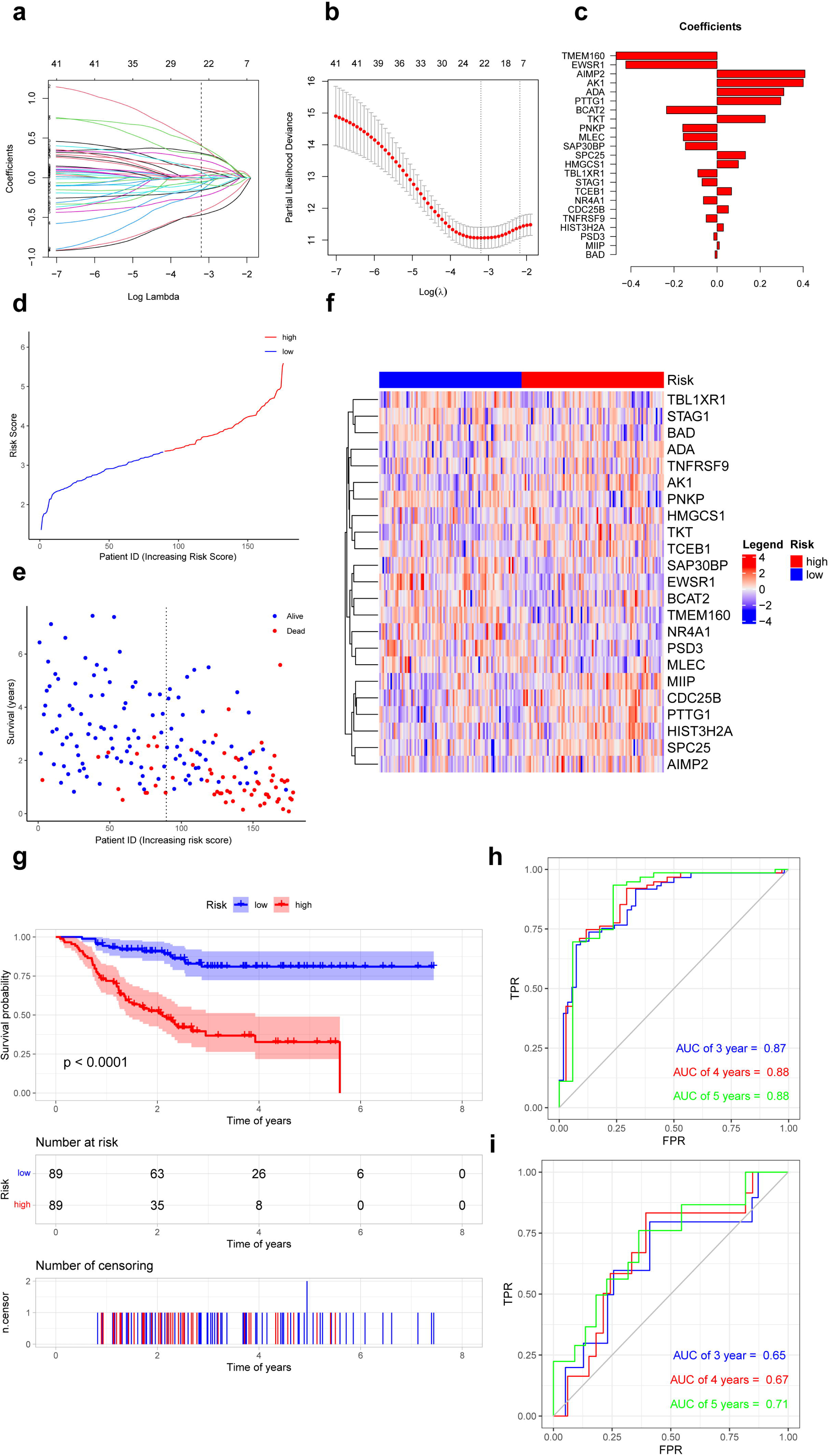
Prognostic differences in triple-negative breast cancer with ferroptosis-related subpopulations. (a) LASSO regression of 23 ferroptosis-related genes. (b) Cross-validation for optimizing the parameter in LASSO regression. (c) Demonstration of regression coefficients corresponding to 23 genes. (d) Graph showing risk scores for all samples. (e) Scatterplot showing recurrence-free survival for all samples. (f) Heat map showing the average expression of 23 genes in the low- and high-risk groups. (g) Kaplan–Meier curves of survival analysis in the low- and high-risk groups. (h) Receiver operating characteristic curves for predicting the recurrence-free survival at 3, 4 and 5 years in training set. (i) Receiver operating characteristic curves for predicting the recurrence-free survival at 3, 4 and 5 years in external validation set.

In external validation set (GSE86166), patients in the high-risk group also had a shorter RFS compared to patients in the low-risk group (p<0.05). This risk model also performed well in predicting RFS in TNBC patients, its AUCs for 3-, 4-, and 5 years RFS were 0.65, 0.67, and 0.71 (Figure 4i).

### 3.5 Analysis of independent prognostic factors

Further, we performed univariate and multivariate Cox analyses to determine whether the risk score could serve as an independent prognostic factor for TNBC patients compared to other common clinicopathologic factors. In the GSE25066 database, Figure 5a shows that both the risk factor and stage were significantly associated with TNBC patients’ RFS and were independent prognostic factors (p<0.05). In the external validation set, Figure 5b shows that only the risk factor can be considered as independent prognostic factor in TNBC patients (p<0.05).

**Fig. 5.**
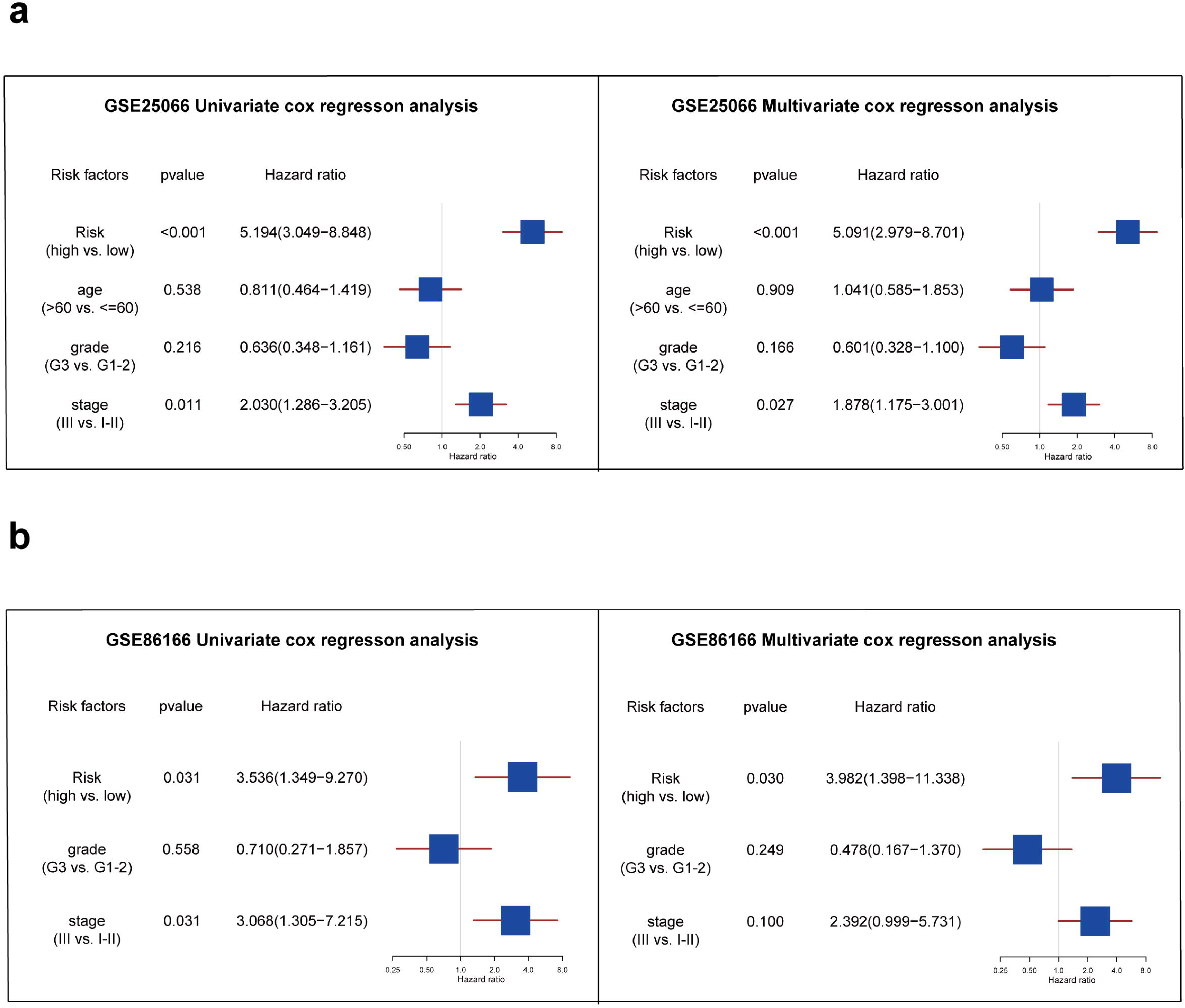
Analysis of independent prognostic factors for triple-negative breast cancer patients. (a) The forest plot showing the results of univariate and multivariate COX regression analysis of risk score, age, grade, and stage in the GSE25066 database. (b) The forest plot showing the results of univariate and multivariate COX regression analysis of risk score, grade, and stage in the GSE86166 database.

### 3.6 Correlation of the risk score with immune microenvironment

Next, we calculated the ImmuneScore, StromalScore, ESTIMATEScore, and TumorPurity for samples in the low- and high-risk groups and compared the differences in scores between the two groups. The results showed that these scores did not differ significantly between the low- and high-risk groups (Figure 6a-f). Then, based on the CIBERSORT algorithm, we found that T cells CD4 memory activated, NK cells resting, and Monocytes had significantly higher proportions in the high-risk group, and the NK cells activated, and mast cells resting had significantly higher proportions in the low-risk group (Figure 6g). Infiltration of these immune cells may play an important role in the clinical course of TNBC patients.

**Fig. 6.**
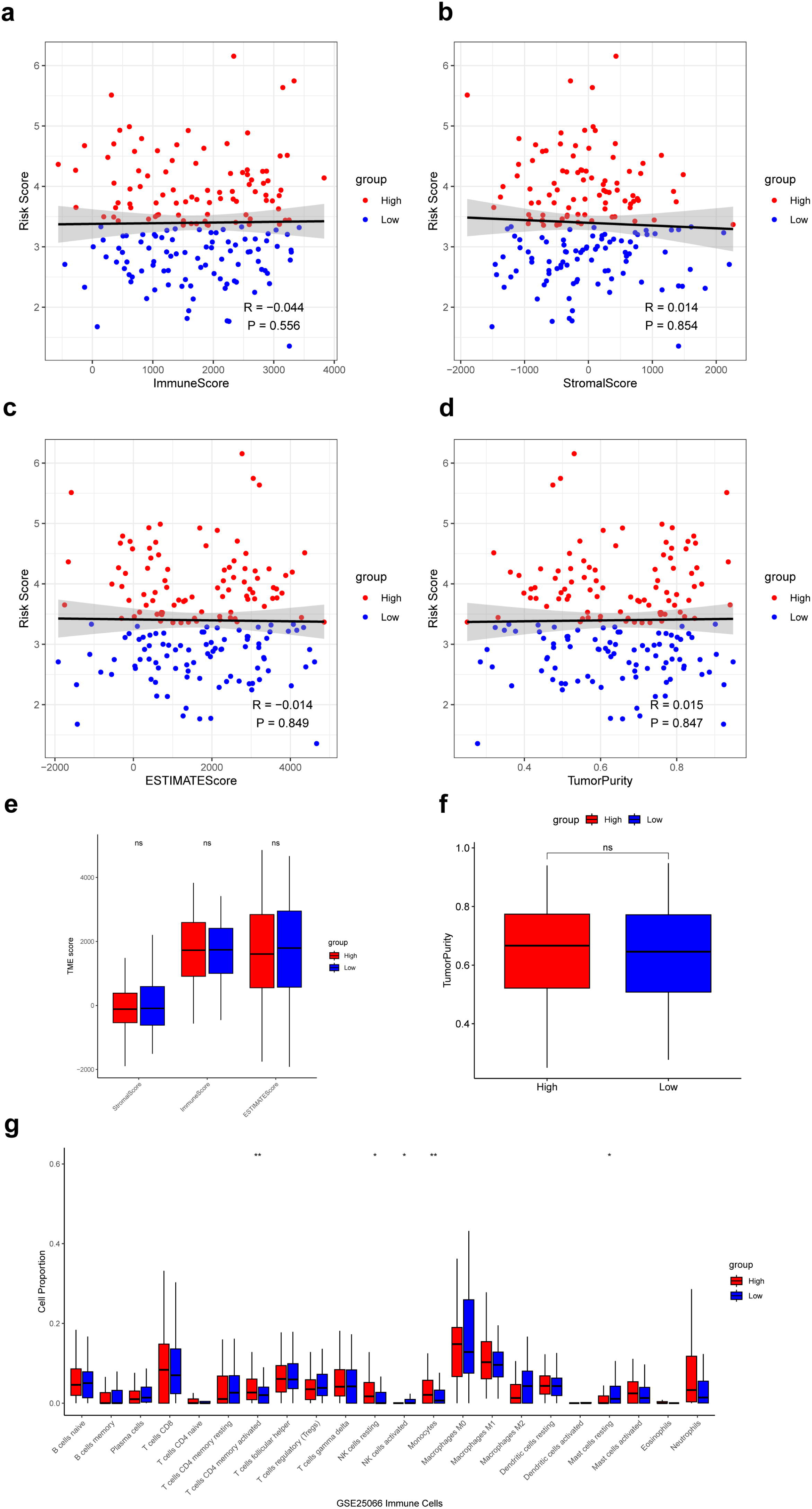
Analysis of immune microenvironment in low- and high-risk groups. (a-d) Correlation of the risk score with the ImmuneScore, StromalScore, ESTIMATEScore, and TumorPurity. (e) Difference of ImmuneScore, StromalScore, and ESTIMATEScore in low- and high-risk groups. (f) Difference of TumorPurity in low- and high-risk groups. (g) Difference of immune infiltration score between low- and high-risk groups calculated by CIBERSORT. * P < 0.05, ** P < 0.01, ns: p>0.05.

### 3.7 Analysis of clinical response to drugs with the risk score

Based on the “pRRophetic” R package, we explored the relationship between the risk score and clinical response to 138 drugs and calculated IC50. As a result, 16 of 50 drugs whose clinical response was significantly associated with the risk score were positively associated with the risk score, and the remaining 34 were negatively associated with the risk score. Further, we found that 27 of drugs negatively associated with the risk score had significantly lower IC50 in the high-risk group, suggesting that patients in the high-risk group may be more sensitive to these drugs, favoring the choice of clinical medication (Figure 7a). And 13 of drugs positively associated with the risk score had significantly higher IC50 in the high-risk group than in the low-risk group, suggesting that patients in the high-risk group may be less sensitive to these drugs (Figure 7b). Finally, we calculated the TIDE scores of patients in the low- and high-risk groups; unfortunately, the TIDE scores were not significantly different between the two groups (Figure 7c).

**Fig. 7.**
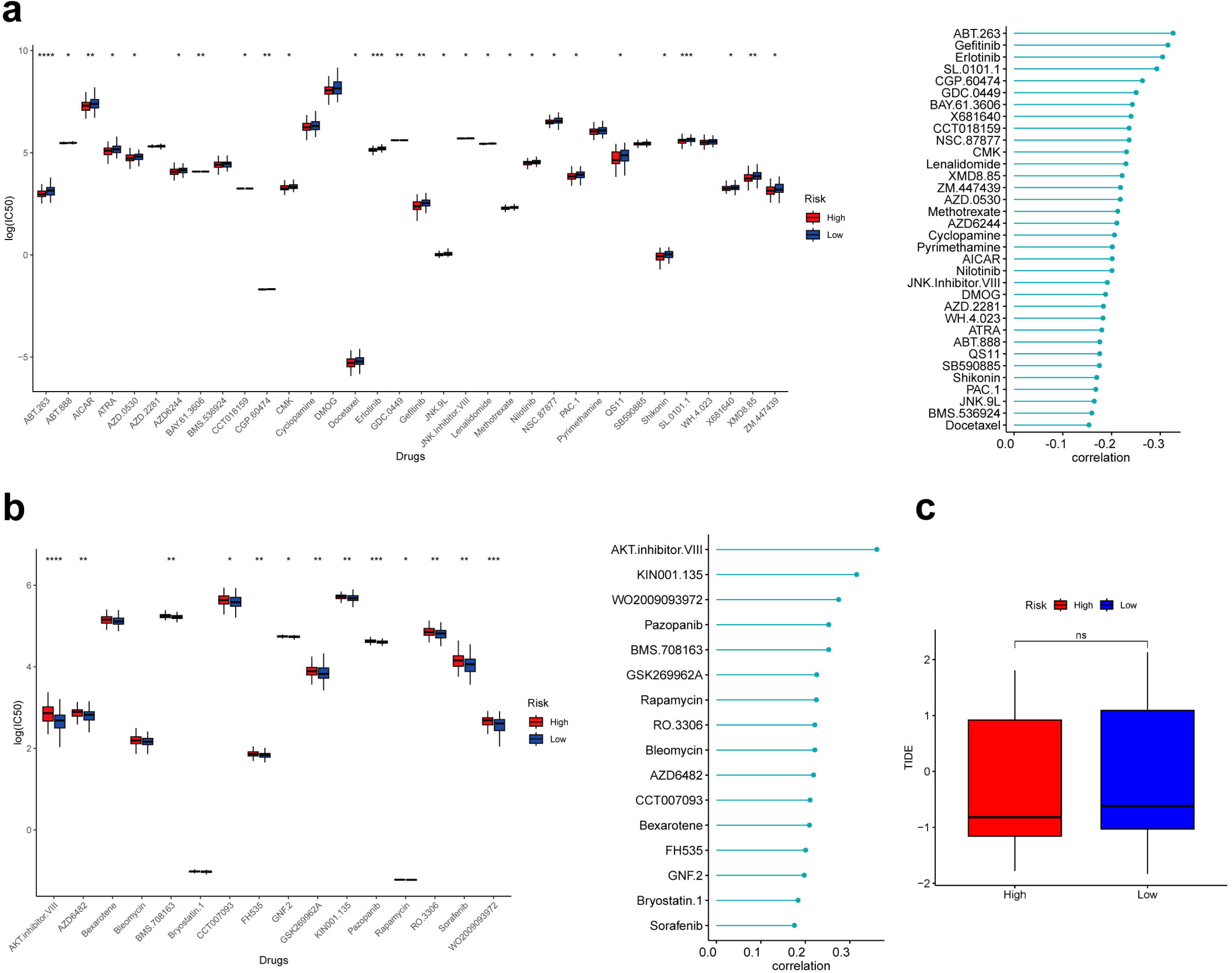
Analysis of immune microenvironment in low- and high-risk groups. (a) Box plot showing differences of IC50 for drugs negatively associated with risk scores in different groups. (b) Box plot showing differences of IC50 for drugs positively associated with risk scores in different groups. (c) Box plot showing differences of TIDE scores in different groups. * P < 0.05, ** P < 0.01, *** P < 0.001, **** P < 0.0001, ns: p>0.05.

## Discussion

TNBC is the most difficult subtype of breast cancer to treat, and survival and efficacy prediction for each individual is a critical step in precision tumor therapy. Several studies have identified ferroptosis-related genes as novel therapeutic targets to enhance treatment efficacy and improve patient prognosis ^[44–47]^. However, to the best of our knowledge, there is a lack of reports on the association of ferroptosis-related genes with prognosis and clinical response to treatment in TNBC. Here, firstly, we revealed ferroptosis-related immune cell clustering in TNBC patients at the single-cell level, demonstrating the ferroptosis heterogeneity in TNBC, which was also supported by other cohorts ^[19]^. On the basis of the above results, we constructed an ferroptosis-related risk score model based on bulk RNA-seq data, which successfully predicted the recurrence-free survival of TNBC patients and validated the robustness of the results in various datasets. This is important for the individualized management of TNBC patients. More importantly, the model can also predict the sensitivity to drugs for TNBC patients with different risk stratification to guide physicians in the selection of drugs for different patients, and it has significant clinical application value.

Understanding the TME in TNBC is essential for improving prognosis and guiding treatment. scRNA-seq enables transcriptomic analysis of individual cells to characterize the cellular diversity in the TME in detail, thereby improving understanding of the disease ^[48–50]^. In this study, our analysis of scRNA-seq data from TNBC revealed that there are multiple immune cells infiltrating in the TME and interacting frequently with each other, and they may impede or promote tumor progression ^[51]^. Since, we found that the average expression of ferroptosis-related genes was significantly different in different T-stages of tumors, which might be strongly correlated with patient prognosis. Further, based on ferroptosis-related genes, a higher proportion of T cells and macrophages in the TME were categorized into different subpopulations to reveal the heterogeneity of ferroptosis-related in TNBC. The T_C2 subpopulation was accompanied by a greater infiltration of NK cells. NK cells have rapid and efficient anti-tumor immunity and their activity is negatively correlated with breast cancer progression ^[52–54]^. The relevant results were validated in the dataset GSE25066. TFs are universal regulators of the transcription of many genes and are closely associated with the progression of cancer ^[55]^. The results indicate that TBX21, which exhibits higher activity in NKT cells, is associated with the expression of SLC7A11, and overexpression of SLC7A11 alters the expression of chemokines, resulting in the infiltration of immune cells such as CD8^+^ T cells and neutrophils ^[56]^. It has been reported that RUNX3 induces ferroptosis by activating the ING1/p53/SLC7A11 signaling pathway, thereby inhibiting the growth of gallbladder cancer ^[57]^. Furthermore, PRDM1 and IRF, which exhibit higher activity in CD8^+^ T cells, induce ferroptosis in cancer cells by inhibiting the transcription of GPX4 and driving the expression of transferrin receptor, respectively, thereby improving the TME ^[58–59]^. FOLR2, SEPP1, MRC1, LYVE1, SLC40A1 and CD163 were highly expressed in the M_C1 subpopulation. Studies have shown that these markers, which are normally present in mammary macrophages from healthy humans and mice, correlate with a favorable prognosis in breast cancer ^[60–62]^. Among them, FOLR2 is positively correlated with antitumor immune players of CD8^+^ T cells, DCs, B cells, and tertiary lymphoid structures, and there is a strong correlation between FOLR2 expression and various immune pathways including T-cell receptor and PD-1 signaling, as well as antigen processing. FOLR2^+^ macrophages are an important component in the initiation of antitumor immunity ^[62]^. In the M_C2 subpopulation, TREM2, FN1, C3, CXCR4, and SPP1 are highly expressed. It has been shown that these markers are lowly expressed in macrophages of healthy breast tissues and highly expressed in tumor tissues ^[62–64]^. Specifically, the absence of TREM2 reshapes macrophage infiltration, fosters the enrichment and activation of T cells and NK cells, and is correlated with an enhanced response to checkpoint blockade therapy ^[63]^. In addition, S100A8 and IFI6 were also highly expressed in the M_C2 subpopulation. Studies have reported that a high percentage of S100A8^+^ myeloid cell infiltration has been suggested as a potential mechanism for worsening the prognosis of TNBC ^[65–66]^. Furthermore, IFI6 is an interferon-stimulated gene with anti-apoptotic and metastasis-promoting effects ^[67]^. These findings demonstrate the heterogeneity of the function of iron death-associated macrophage subpopulations, which may play different roles in the anti-tumor process. In the original literature, Prabhakaran S et al. ^[35]^ classified breast cancer into nine clusters (Ecotypes) based on single-cell characteristics, which exhibited distinct cellular compositions and clinical outcomes. Among them, Ecotypes-2 is primarily composed of LumA and Normal-like tumors, and patients in this group have the best prognosis, whereas Ecotype-3 is enriched with Basal_SC, Cycling, Luminal_Progenitor, and a Basal bulk PAM50 subtype, and patients in this group have the worst prognosis. In comparison, this study focuses on TNBC, which has a poorer prognosis. The different distributions of T cell and macrophage subtypes in different TNBC patients the study results revealed that ferroptosis-related TME in TNBC patients may be valuable for patient survival and outcome prediction. In the future, altering the degree of macrophage or T-cell infiltration in different subpopulations could be potentially valuable in improving the prognosis of TNBC patients.

Intratumor heterogeneity is one of the most important factors contributing to poor clinical outcomes in breast cancer, and ferroptosis is one of the major pathways of activation in highly heterogeneous tumors ^[68–70]^. In order to better address the problem of accurately predicting the prognosis of TNBC, we firstly performed the correlation analysis of ferroptosis-related genes with RFS and constructed a risk score model in the publicly available dataset of bulk RNA-seq. TNBC patients were categorized into low- and high-risk groups based on the risk score. In the high-risk group, TNBC patients had higher expression levels of the MIIP, HIST3H2A, CDC25B, TCEB1, HMGCS1, SPC25, TKT, PTTG1, ADA, AK1, and AIMP2 genes. Among them, the cell division cycle 25 (CDC25) family of proteins is dual-specific tyrosine phosphatases responsible including three subtypes, CDC25A, CDC25B and CDC25C, which are used to regulate cell cycle transitions ^[71–72]^. CDC25B dephosphorylates and activates cyclin-dependent kinase/cyclin complex (CDK1/Cyclin B), which have been shown to be overexpressed in breast cancer and promote cell cycle progression ^[72–73]^. Spindle component 25 (SPC25) is a key component of the nuclear division cycle 80 (NDC80) complex, and its high expression promotes tumor cell proliferation by inducing mitotic disorder ^[74]^. Study shows that SPC25 promotes the proliferation of breast cancer cells, and high levels of SPC25 mRNA levels are associated with high recurrence rates and low survival rates in breast cancer patients ^[75]^. Pituitary tumor transforming gene 1 (PTTG1) has been demonstrated to promote proliferation, migration and invasion of cancer cells and is a gene that promotes breast cancer development ^[76]^. At the mRNA level, PTTG1 is more present in patients with high histologic grade and with lymph node metastasis, is significantly associated with low patient survival, and is a biomarker suggestive of poor prognosis in breast cancer, which may be related to the regulation of the cell cycle by PTTG1 to promote the development of breast cancer cells ^[77]^. In the low-risk group, TNBC patients had higher expression levels of the TMEM160, EWSR1, BCAT2, PNKP, MLEC, SAP30BP, TBL1XR1, STAG1, NR4A1, TNFRSF9, PSD3, and BAD genes. These genes have different roles in regulating the TME in colorectal cancers, bladder cancers, lymphomas, etc., but they have been rarely reported in TNBC ^[78–80]^. In future studies, these genes are valuable to investigate in the regulation of anti-TNBC.

In TNBC, we found that the risk score model we constructed had independent predictive power for RFS and the predictive performance of this score was stable in an external validation set. Considering the impact of the tumor immune microenvironment on the prognosis of TNBC patients, we further explored the differences in immune cell infiltration in different subgroups of patients. The results showed that T cells CD4 memory activated, NK cells resting, and Monocytes had higher proportions in the high-risk group. This is in line with previous studies, such as that CXCL7 expressed and released by monocytes can stimulate cancer cell migration, invasion and metastasis ^[81]^. While in the low-risk group, the proportion of NK cells activated and mast cells resting was higher. It has been suggested that the effect of mast cells on tumor invasiveness may vary among different breast cancer subtypes ^[82–83]^. Our findings suggest that infiltration of mast cells resting in the TME is more favorable to the prognosis of patients.

For high-risk TNBC patients, it is crucial to seek more effective drugs for adjuvant therapy in clinical practice. In our study, we found that ABT-263 (navitoclax) and erlotinib exhibited the greatest differences in drug sensitivity among different patient groups. ABT-263 is a mimetic of B-cell lymphoma-2 (BCL-2) homology 3 (BH3) that can increase cellular reactive oxygen species (ROS) levels and induce TNBC cell apoptosis by inhibiting the function of anti-apoptotic proteins (BCL-2, BCL-XL, and BCL-W) ^[84–85]^. ROS is a key regulatory factor in the occurrence of ferroptosis, and excessive production of ROS can lead to the accumulation of lipid peroxides, ultimately triggering cellular ferroptosis ^[86–87]^. In addition, erlotinib is an epidermal growth factor receptor (EGFR) tyrosine kinase inhibitor (TKI) that exerts its effects by competitively binding to the ATP site within the catalytic domain and inhibiting the phosphorylation of EGFR, thereby improving the progression-free survival of patients with EGFR mutations ^[88–89]^. Studies have shown that β-Elemene can enhance the sensitivity of EGFR-mutated non-small cell lung cancer to erlotinib by upregulating lncRNA H19 and inducing ferroptosis, providing a new approach to overcoming drug resistance in TNBC ^[90]^. EGFR is a receptor tyrosine kinase that promotes the proliferation and invasion of breast cancer by stimulating multiple oncogenic pathways such as Ras-Raf-MEK-ERK, PI3K-AKT-mTOR, and Src-STAT3 ^[91]^. The high sensitivity of high-risk patients to erlotinib may be attributed to the higher expression of EGFR in their tumors. Currently, the treatment of TNBC mainly relies on systemic chemotherapy, while small molecule inhibitors and targeted drugs induce cell death through various pathways, holding great promise for anti-tumor treatment in high-risk patient groups.

This study has some limitations. First, this study is based on the analysis of transcriptomic data, and the integration of multi-omics data is needed to improve the predictive performance of the model. Second, this study is a retrospective analysis of a public dataset, and future in vivo, in vitro experiments as well as prospective clinical trials are needed to further validate the research findings.

## Conclusions

In conclusion, this study revealed the TNBC ferroptosis-mediated tumor immune cell clustering based on scRNA-seq data, and the degree of infiltration of different subpopulations of cells had different roles in the prognosis of TNBC patients. On this basis, combined with bulk RNA-seq data, the survival prediction model for TNBC patients was established with excellent predictive performance and stability. The high-risk TNBC patients screened by this prediction model and their sensitive therapeutic drugs can help to guide physicians in disease monitoring and precise treatment. The related genes screened also provide important value for TNBC patients to find potential therapeutic targets to improve their prognosis.

## Acknowledgements

Our thanks go to the Gene Expression Omnibus (GEO) database and FerrDb for providing essential data resources. We extend our appreciation to the participating institutions and our dedicated colleagues for their contributions and expertise. Special recognition is given to the patients and clinical staff for their invaluable participation and assistance. We also thank the reviewers for their insightful feedback.

## List of abbreviations

TNBC: Triple-negative breast cancer
TME: tumor microenvironment
RNA-seq: RNA sequencing
scRNA-seq: single-cell RNA-seq
AUCs: Area Under the Curves
GEO: Gene Expression Omnibus
TCGA: The Cancer Genome Atlas
QC: quality control
HVGs: highly variable genes
PCA: principal component analysis
t-SNE: t-distributed stochastic neighbor embedding
NMF: Non-negative matrix decomposition
PCBC: Progenitor Cell Biology Consortium
OCLR: one-class logistic regression
SCENIC: Single-cell Regulatory Network Inference and Clustering tool
TF: transcription factor
RFS: recurrence-free survival
LASSO: Least Absolute Shrinkage and Selection Operator
ROC: receiver operating characteristic
TIDE: tumor immune dysfunction and exclusion
CAFs: Cancer-associated Fibroblasts
PVL: Perivascular-like Cells
DCs: Dendritic Cells
LASSO: Least Absolute Shrinkage and Selection Operator
CDC25: cell division cycle 25
CDK1/Cyclin B: cyclin-dependent kinase/cyclin complex
SPC25: Spindle component 25
NDC80: nuclear division cycle 80
PTTG1: Pituitary tumor transforming gene 1
BCL-2: B-cell lymphoma-2
BH3: B-cell lymphoma-2 homology 3
ROS: Reactive oxygen species
EGFR: Epidermal growth factor receptor
TKI: Tyrosine kinase inhibitor

## Declarations

## Ethics approval and consent to participate

Not applicable

## Consent for publication

Not applicable

## Availability of datasets

The data used in this study are available in the public datasets. Nine single-cell RNA-seq TNBC samples were obtained from the Gene Expression Omnibus (GEO) database (https://www.ncbi.nlm.nih.gov/geo/query/acc.cgi). The data for external validation were obtained from the GEO database. The data of 471 genes associated with ferroptosis were collected from the FerrDb website (http://www.zhounan.org/ferrdb/current/).

## Authors’ contributions

YW: Conceptualization, Methodology, Formal analysis, Investigation, Writing-original draft preparation, Writing-review & editing, Project administration and Funding acquisition. XTG: Conceptualization, Methodology, Formal analysis, Investigation, Writing-original draft preparation, Writing-review & editing, Resources and Data curation. LSG: Conceptualization, Methodology, Formal analysis, Investigation, Writing-original draft preparation, Writing-review & editing, Resources and Data curation. DY: Writing-original draft preparation and Writing-review & editing. YH: Writing-original draft preparation and Writing-review & editing. QL: Conceptualization, Methodology, Formal analysis, Investigation, Writing-original draft preparation, Writing-review & editing, Visualization and Supervision. HQ: Conceptualization, Methodology, Formal analysis, Investigation, Writing-original draft preparation, Writing-review & editing, Visualization and Supervision. All authors: reading and approval of the final manuscript.

## Additional files File

Appendix 1-figure 1.TIF. Appendix 1-figure 2.TIF. Appendix 1-figure 3_revised.TIF. Appendix 1-figure 4_revised.TIF. Appendix 1-figure 5.TIF. Appendix 1-figure 6.TIF. Appendix 1-figure 7.TIF. Appendix 1-Supplementary Figure 1.TIF. Appendix 1-Supplementary Figure 2.TIF. Appendix 1-Supplementary Figure 3.TIF. Figure 1–figure supplement.pdf. Figure 2–figure supplement.pdf. Figure 3–figure supplement.pdf. Figure 4–figure supplement.pdf. Figure 5–figure supplement.pdf. Figure 6–figure supplement.pdf. Figure 7–figure supplement.pdf. Supplementary Figure 1-figure supplement. Supplementary Figure 2-figure supplement. Supplementary Figure 3-figure supplement. Appendix 2-ICMJE.zip. Appendix 3-collaborators.xlsx. Supplementary Table 1. Supplementary Table 2.

---

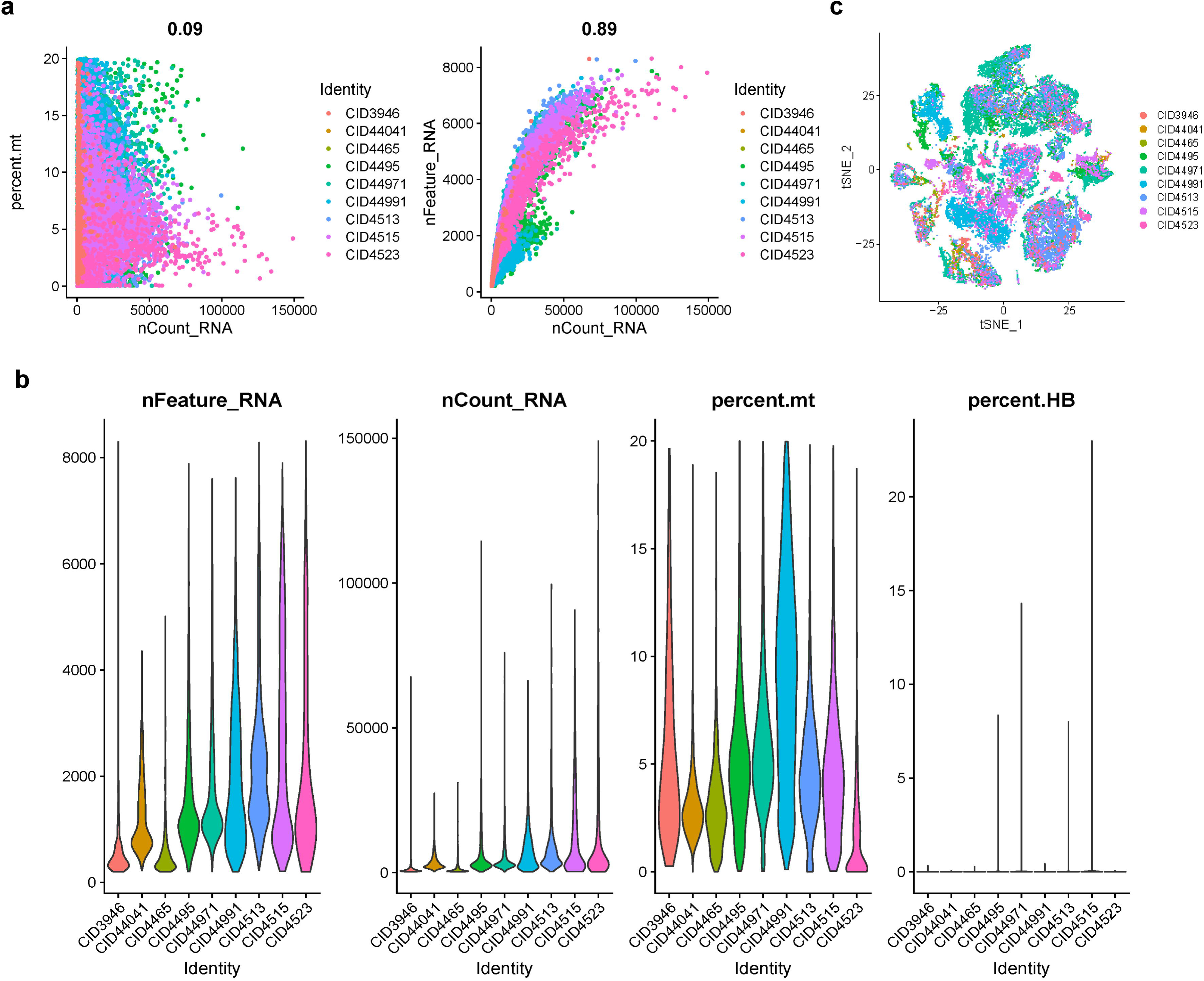

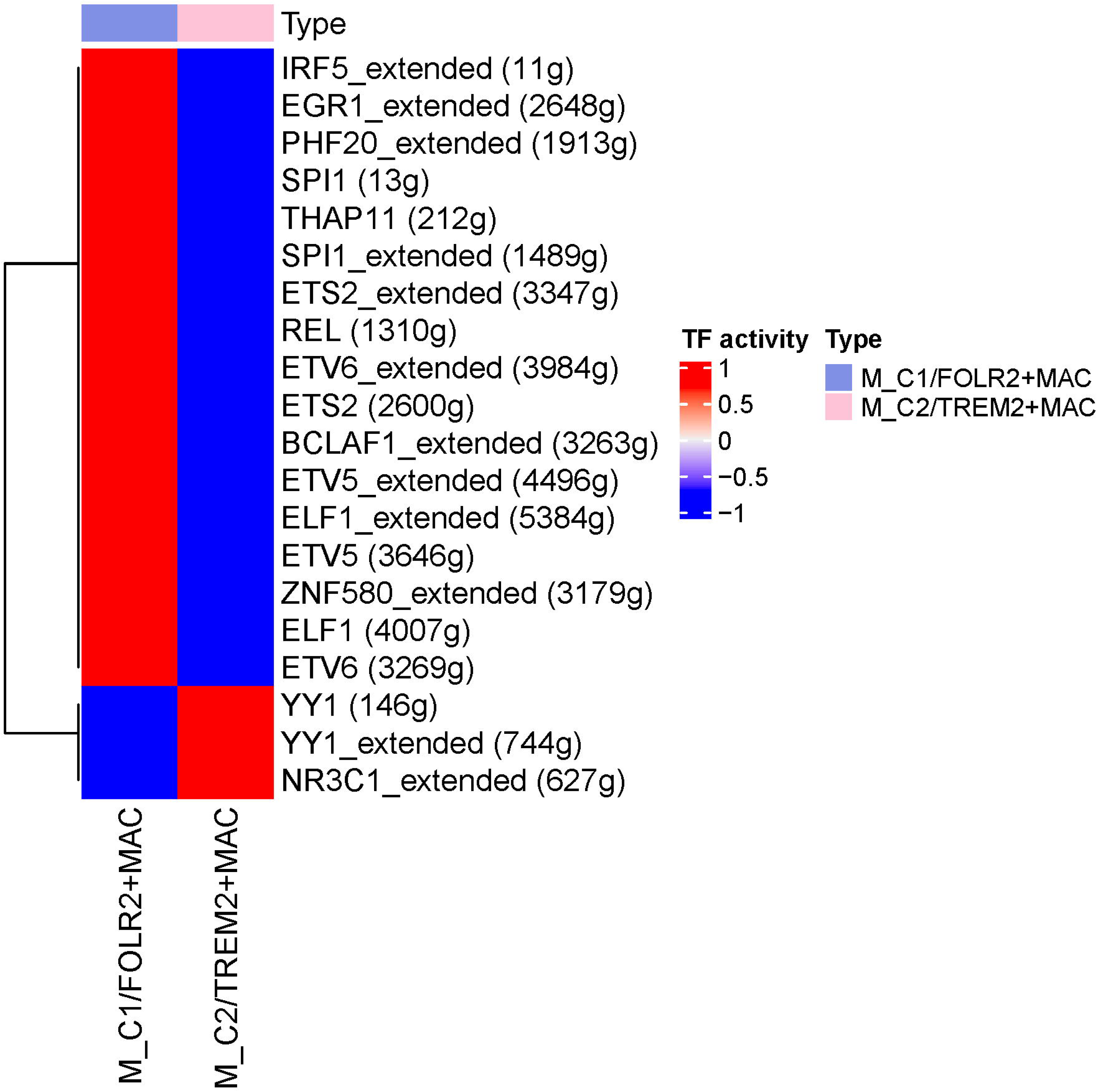

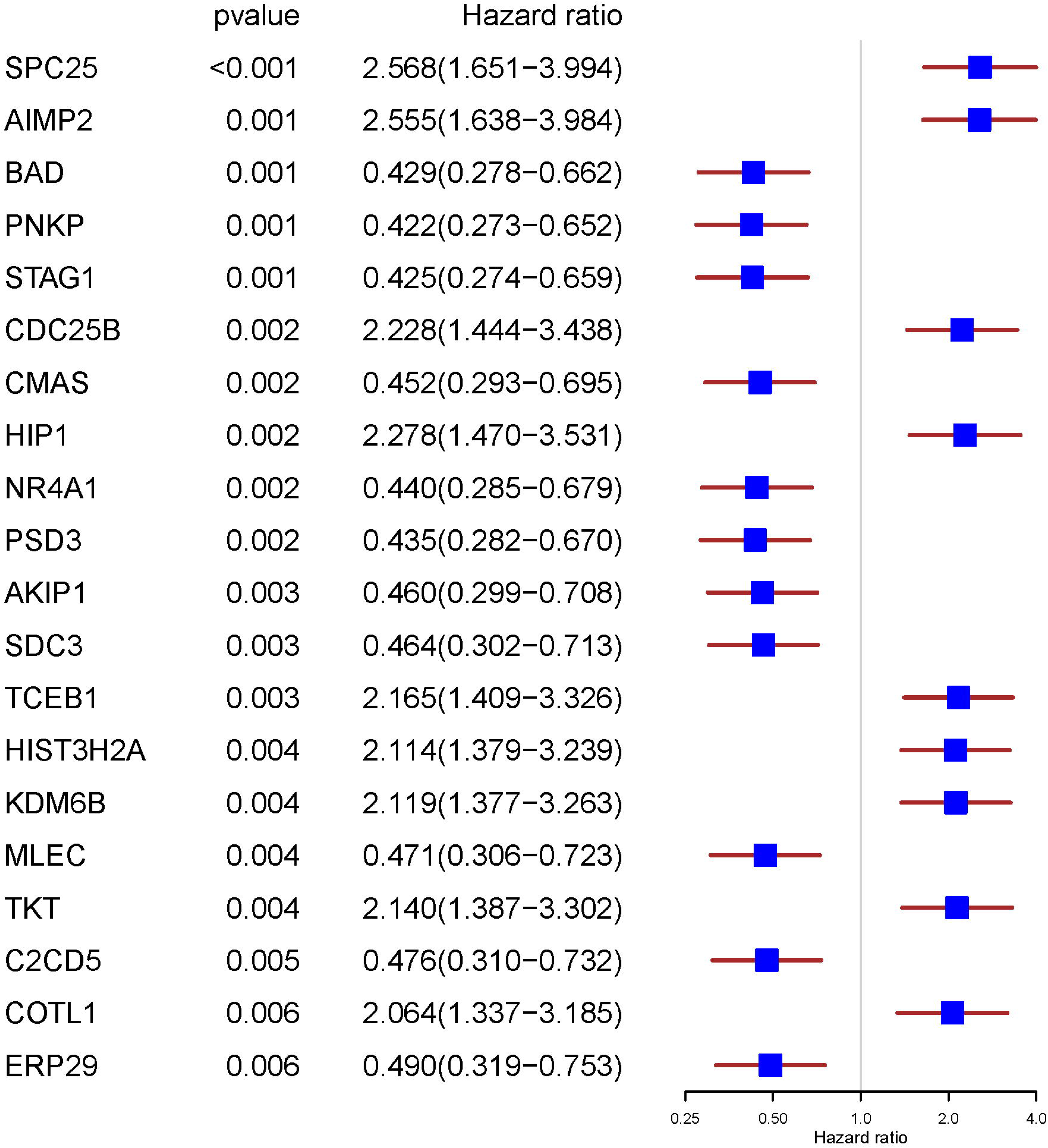

