## Supplementary material for "Integrated Analysis of Single-Cell and Bulk RNA-Seq Data reveals that Ferroptosis-Related Genes Mediated the Tumor Microenvironment predicts Prognosis, and guides Drug Selection in Triple-Negative Breast Cancer": Legend of Supplementary Fig 1

*Supplementary Fig. 1 Single-cell RNA-seq data preprocessing in triple-negative breast cancer. (a) t-SNE plot of the 9 major cell clusters. (b) t-SNE plot of the 29 minor cell clusters. (c) t-SNE plot of the 12 major cell clusters.*
